# Regulatory and metabolic integration accompany plasmid domestication during adaptation to a plant-associated lifestyle

**DOI:** 10.64898/2026.09.30.755635

**Authors:** Shreya Vichare, Kai Mahn-Ripcke, Devani Romero Picazo, Ryszard Soluch, Nils Hülter, Tal Dagan

**Author notes:** Corresponding author: Tal Dagan. These authors contributed equally (Shreya Vichare, Kai Mahn-Ripcke).

## Abstract

Plasmids are major drivers of evolutionary innovation and ecological adaptation in prokaryotes. Plasmids that confer beneficial functions on their host may become domesticated through long-term coevolution, resulting in stable vertical inheritance within a bacterial lineage. How this process contributes to the evolution of bacteria associated with eukaryotic hosts remains poorly understood. Here, we investigate the function of the two large domesticated plasmids, LPP1 and LPP2, in *Pantoea agglomerans* associated with wheat. We show that loss of LPP1 reduces bacterial fitness during wheat colonization and impairs the utilization of diverse carbon sources. Comparative transcriptomics further reveal that curing of either plasmid extensively remodels chromosomal gene expression under both laboratory and wheat-associated conditions, whereas genome-scale metabolic reconstruction predicts that many metabolic pathways are jointly encoded by plasmid and chromosome. This functional integration extends beyond plasmid-specific pathways, as plasmid curing disrupts the utilization of substrates metabolized exclusively by chromosomally-encoded enzymes. Our results indicate that LPP1 promotes *P. agglomerans* association with wheat primarily through its regulatory and metabolic integration with the chromosome rather than through the simple addition of accessory genes. We propose that LPP1, and likely LPP2, represent chromosome-like symbiosis plasmids whose domestication reflects long-term selection for plant-associated lifestyle in *Pantoea*. Our findings suggest that persistent selection imposed by symbiotic lifestyles drives plasmid domestication, progressively transforming mobile plasmids into chromosome-like replicons that are integral components of bacterial genomes.

## Introduction

Plasmids are autonomously replicating genetic elements that are ubiquitous in prokaryotic cells. Their acquisition may confer novel functions on the host through plasmid-borne genes. Beneficial plasmid-encoded traits include metabolic capabilities, such as the utilization of specific carbon sources ^1^, as well as functions that facilitate the exploitation of new ecological niches, such as motility, adhesion and biofilm formation ^2–4^. Plasmids are commonly regarded as accessory genetic elements whose presence is transient and restricted to particular strains, depending on environmental conditions and historical contingency. A canonical example is provided by mobile antibiotic-resistance plasmids, which are sporadically acquired by members of the *Enterobacteriaceae* associated with humans, and whose persistence or loss depends on the selective pressure imposed by antibiotics ^5,6^. Coevolution between plasmids and their hosts may give rise to stably inherited, non-mobile plasmids whose persistence is independent of environmental conditions, similar to chromosomes ^7^. The emergence of such ‘domesticated’ plasmids (also termed chromids or secondary chromosomes) has been hypothesized to coincide with bacterial adaptation to novel ecological niches that is facilitated by the plasmid-encoded traits ^8–10^.

Adaptation to eukaryotic hosts provides a prime example of a major ecological transition in bacterial evolution ^11,12^. Traits that promote a host-associated lifestyle range from functions involved in the exchange of resources with the host, such as nutrients or protective compounds, to mechanisms that facilitate colonization and persistence within host habitats ^13,14^. Examples of plasmid-borne functions involved in resource exchange include nitrogen fixation in *Rhizobia* associated with legumes ^15,16^, essential amino acid biosynthesis in *Buchnera* spp., the obligate symbionts of aphids ^17,18^, vitamin B12 biosynthesis in *Dinoroseobacter shibae*, a symbiont of the dinoflagellate *Prorocentrum minimum* ^19,20^ and vitamin B5 in the Riesia sp. symbiont of human lice ^21,22^. Plasmid-borne mechanisms that facilitate host colonization include fimbria adhesion in enterohemorrhagic *Escherichia coli* colonizing the human gut ^2,23^, as well nodulation or tumor induction in plant-associated *Rhizobia* and *Agrobacterium* spp. ^24^. Notably, the male-killing factor that enhances transmission of *Spiroplasma poulsonii* through Drosophila hosts is encoded on a plasmid ^25,26^. Plasmids carrying traits that are beneficial or essential for host association are termed symbiosis plasmids.

Members of the genus *Pantoea* are widely associated with plants and include both plant pathogens and commensals ^27^. Plant-associated *Pantoea* species harbor the large *Pantoea* plasmid LPP1, which exhibits hallmarks of plasmid domestication and is thought to have contributed to the evolution of an endophytic lifestyle in *Pantoea* ^28^. Several taxa carry an additional plasmid, LPP2, which is considered domesticated as well. Phylogenomic analyses indicate that both plasmids have an ancient origin within the genus and have evolved predominantly through vertical inheritance. Moreover, replication of LPP1 is coordinated with that of the host chromosome, a defining feature of plasmid domestication ^29^. LPP1 homologs encode multiple traits considered beneficial for an endophytic lifestyle. For example, curing of LPP1 in the maize pathogen *P. ananatis* strain DZ-12 (termed pDZ-12) reduces biofilm formation and disease symptoms in infected maize plants ^30^. Likewise, curing of LPP1 in *P. vagans* reduces the ability to utilize several carbon sources, including maltose and salicin, and decreases swarming motility and indole production ^31,32^. LPP2 homologs (also termed pAggl) have been predicted to contribute to sucrose utilization ^33,34^. However, whether LPP1 and LPP2 directly contribute to the *Pantoea* plant-associated lifestyle remains unresolved.

Here, we investigate the functions of LPP1 and LPP2 in *Pantoea agglomerans* strain R1, which we previously isolated from wheat grains ^35^. Several *P. agglomerans* strains are recognized plant pathogens, and their ability to induce necrosis or tumor formation has been linked to the presence of virulence plasmids ^34,36–39^. Nevertheless, *P. agglomerans* is a common member of the grain microbiome of multiple cereal species, consistent with a commensal association with cereals and other plants ^40–43^. Moreover, a recent study demonstrated vertical inheritance of *P. agglomerans* in wheat ^44^. By generating plasmid-cured derivatives of strain R1, we investigated the contributions of LPP1 and LPP2 to wheat seedling colonization, gene expression, and metabolic capacity. We show that LPP1 promotes root colonization of wheat seedlings and identify specific metabolic functions contributed by both plasmids. Furthermore, our results reveal extensive transcriptional and metabolic integration of LPP1 and LPP2 with the chromosome of *P. agglomerans* R1.

## Results

### Plasmid LPP1 is beneficial for *P. agglomerans* R1 colonization of wheat

The plasmids LPP1 and LPP2 in *P. agglomerans* R1 comprise 529 and 145 genes, respectively, and are similar in size to their homologs in other plant-associated *P. agglomerans* strains ^28,29^. Plasmid LPP1 is primarily composed of metabolic genes, including genes for the transport and metabolism of amino acids, carbohydrates and inorganic ions. The LPP2 gene functions are mostly within cellular processes, including cell wall and membrane biogenesis as well as signal transduction. Both plasmids comprise multiple genes annotated as transcriptional regulators (Fig. 1A; COG: K). To study the function of plasmids LPP1 and LPP2 in *P. agglomerans* we generated plasmid cured derivatives of *P. agglomerans* strain R1 (hereafter Pantoea), which we previously isolated from wheat grains ^35^. The resulting strains carry a single plasmid only -either LPP1 (termed ΔLPP2) or LPP2 (termed ΔLPP1) - or no plasmid at all (termed ΔLPP1&2) (Fig. 1B). Whole-genome sequencing of the generated plasmid-cured strains revealed several chromosomal variants relative to the wild type but no large-scale deletions (Table S1). Competition experiments between the plasmid-cured strains and the wild type under *in vitro* conditions revealed a negative fitness effect of plasmid loss (Fig. 1C; Fig. S1A). The similar fitness reduction observed for strains lacking either a single plasmid or both plasmids suggest that simultaneous curing of LPP1 and LPP2 does not have a synergistic negative fitness effect under *in vitro* conditions.

**Figure 1.**
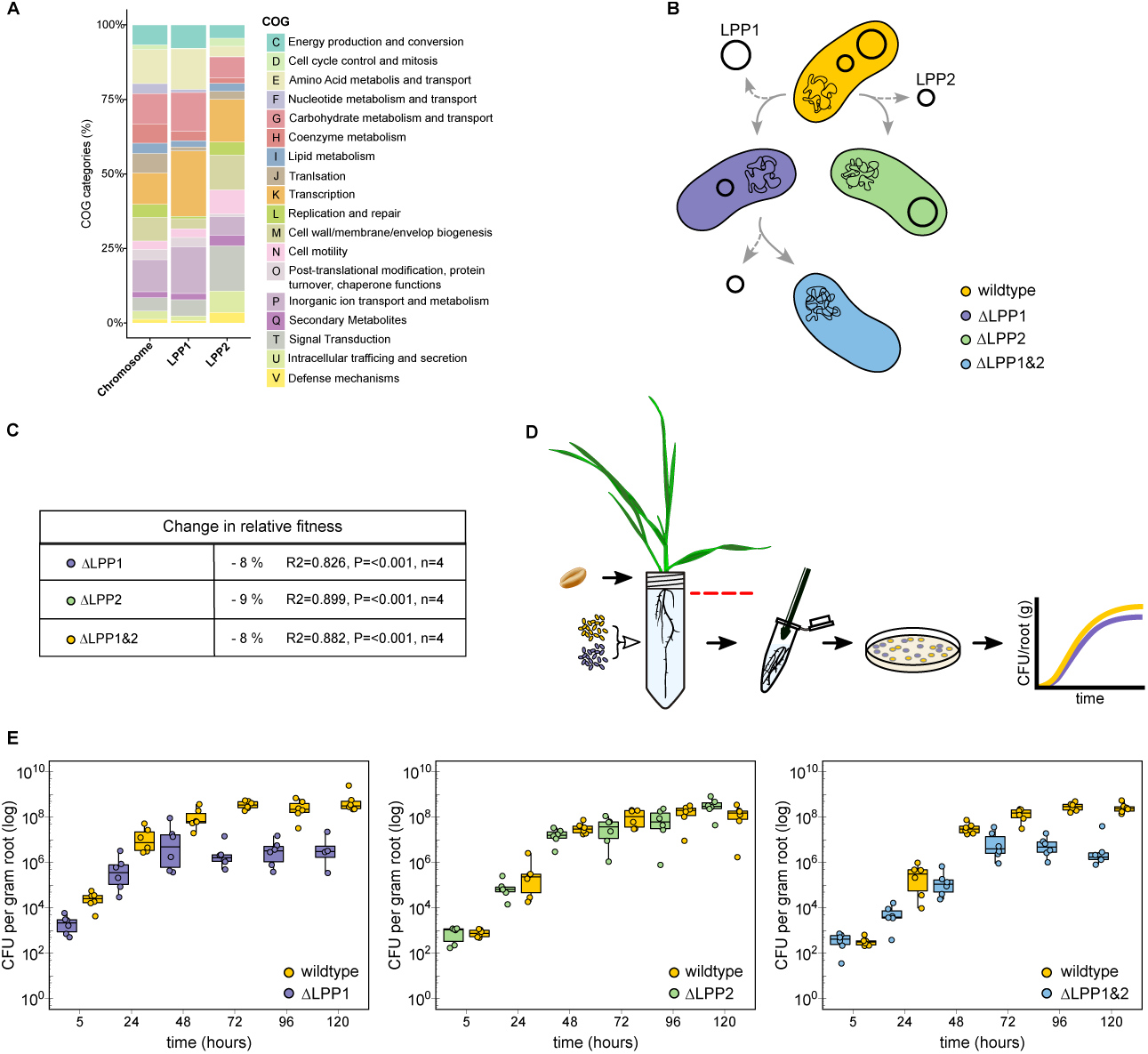
The LPP1 is beneficial for Pantoea colonization of wheat roots. **(A)** Functional composition chromosome, LPP1, and LPP2 genes classified by COG categories. **(B)** Schematic of the wild-type *P. agglomerans* R1 and plasmid-cured derivatives carrying LPP1 only (ΔLPP2), LPP2 only (ΔLPP1), or neither plasmid (ΔLPP1&2). The ΔLPP1 strain was obtained by isolating a spontaneous LPP1-free segregant, whereas LPP2 was cured from the wild type and ΔLPP1 using a non-replicative gene-targeting vector, generating the ΔLPP2 and ΔLPP1&2 strains. The established strains do not show any major deletion in their chromosome (Table S1). **(C)** Relative fitness of the plasmid-cured strains compared with the wild type during *in vitro* competition (n = 4). See detailed results in Fig. S1. **(D)** Schematic of the pairwise competition assay in sterilized wheat seedlings. Individual seedlings were inoculated with mixtures of the wild type and a plasmid-cured strain. Root-associated bacterial populations were quantified by plating homogenized root systems of individual plants. **(E)** Population dynamics of the wild type and plasmid-cured strains during pairwise competition in wheat roots over five days. Points represent colony-forming units (CFUs) recovered from individual root systems; boxplots show the median and interquartile range, with whiskers indicating the data range (n = 6). Similar population dynamics were observed in the plant media (Fig. S1). In the absence of competition, plasmid-cured strains reached similar root population sizes as the wild type (Fig. S1).

To test the contribution of LPP1 and LPP2 to Pantoea fitness during wheat colonization, we performed *in planta* competition experiments between the wild type and each of the plasmid-cured strains. Germ-free wheat seedlings were inoculated at an early developmental stage with a mixture of the two competing strains. The population density of each strain associated with wheat roots, including both the root surface and endosphere, was determined by colony-forming unit (CFU) counts from plated root homogenates (Fig. 1D). The ΔLPP1 and wild-type strains initially exhibited similar population dynamics *in planta*. However, after 72 h, ΔLPP1 only reached a density of approximately 5 × 10^6 CFU/g root, compared with 2 × 10^8 CFU/g root for the wild type. By the end of the experiment, wild-type density was approximately 100-fold higher than that of ΔLPP1 (Fig. 1E). The relative frequencies of the competing strains in the plant medium showed a similar pattern (Fig. S1; Table S2). These differences corresponded to a significant fitness advantage of the wild type over ΔLPP1 *in planta* (using linear regression of the ratio ΔLPP1/wild type abundance over time, b = 0.54, P = 0.0001).

In contrast, ΔLPP2 and the wild type exhibited similar population dynamics and reached comparable densities after 72 h, which remained stable until the end of the experiment (Fig. 1E). Accordingly, the ΔLPP2-to-wild-type ratio did not change significantly over time (using linear regression of the ratio ΔLPP2/wild type abundance over time, b = −0.17, P = 0.36), indicating no detectable fitness difference between the strains under these conditions. Competition between ΔLPP1&2 and the wild type yielded an outcome similar to that observed for ΔLPP1. After 72 h, ΔLPP1&2 reached a significantly lower density in the roots than the wild type, and by the end of the experiment, wild-type density was approximately 100-fold higher than that of ΔLPP1&2 (Fig. 1E; using linear regression of the ratio ΔLPP1&2/wild type abundance over time, b = 0.56, P = 0.0089). Taken together, these results show that LPP1 confers Pantoea a fitness advantage during wheat-root colonization at early stages of plant development, whereas LPP2 has no detectable effect under the conditions tested.

### Plasmid genes involved in metabolism and cellular processes are highly expressed *in vitro* and *in planta*

To investigate the contribution of plasmid genes to Pantoea traits, we examined gene expression in the wild type and plasmid-cured strains under *in vitro* and *in planta* conditions using differential transcriptomics (Fig. 2A). Gene expression levels in the wild type were significantly positively correlated between the two conditions for all three replicons; with approximately 50% of the genes showing no significant difference in expression level between conditions (Fig. 2B).

**Figure 2.**
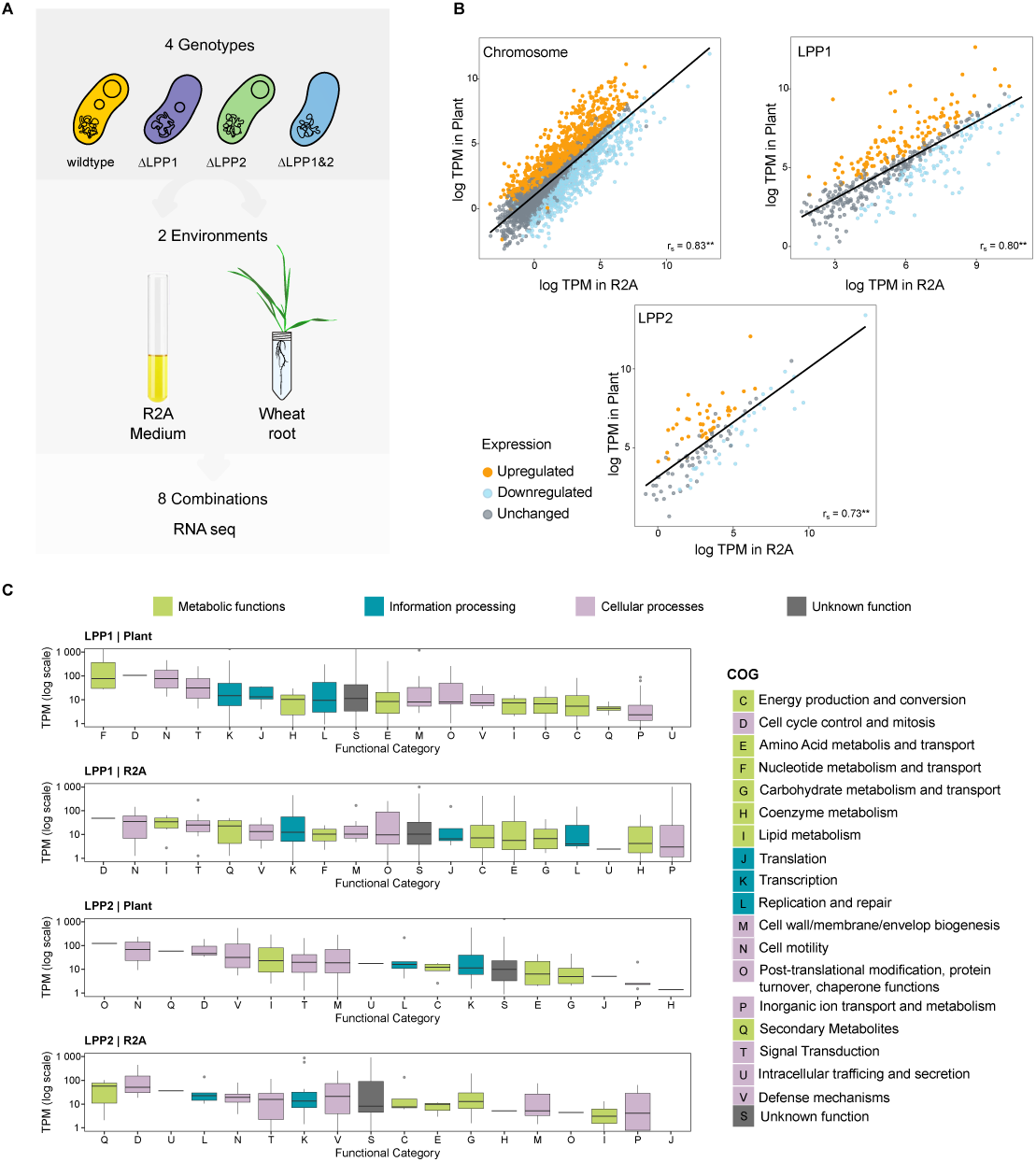
Plasmid gene expression patterns *in planta* and *in vitro*. **(A)** Experimental design for RNA-seq analysis of the wild-type and plasmid-cured *P. agglomerans* R1 derivatives grown in R2A medium or associated with wheat roots. **(B)** Scatter plots of log-transformed transcripts per million (TPM) values for chromosome-, LPP1-, and LPP2-encoded genes in the wild type grown in R2A medium (x axis) or associated with wheat roots (y axis). Each point represents a gene. Genes exhibiting significant differential expression between conditions, as determined by DESeq2 (*P* < 0.05), are colored according to expression change, whereas genes without significant differential expression are shown as unchanged. The level of gene expression is highly positively correlated between the two conditions (** stands for P < 0.01). **(C)** Distribution of gene expression levels across COG functional categories for the LPP1, and LPP2 under *in vitro* and *in planta* conditions. Boxplots show the median and interquartile range, with whiskers indicating the data range. TPM, transcripts per million.

The median expression levels of LPP1 and LPP2 genes varied across functional categories and cultivation conditions, with the highest values generally observed for genes involved in metabolism and cellular processes (Fig. 2C). Among LPP1 genes, categories associated with motility (N), cell cycle control (D), and signal transduction (T) exhibited the highest median expression levels under both conditions. Genes involved in nucleotide transport and metabolism (F) showed the highest median expression *in planta* but ranked lower *in vitro*. Transcriptional regulators (K) encoded on LPP1 were highly expressed under both conditions and showed higher expression *in planta*. Among LPP2 genes, the highest median expression levels were observed for categories related to secondary metabolite biosynthesis (Q) and cell cycle control (D). In contrast, genes involved in protein turnover and degradation (O) were among the most highly expressed under *in vitro* conditions but among the least expressed *in planta*. Additionally, genes associated with inorganic ion transport and metabolism (P) exhibited the lowest median expression levels for both plasmids (Fig. 2C).

### Differentially expressed plasmid genes encode traits associated with endophytic lifestyle

Differential transcriptomic analysis of the wild type strain identified 115 LPP1 genes (21%) that were significantly overexpressed *in vitro* and 131 genes (24%) that were significantly overexpressed *in planta* (Fig. 2B). Here we highlight several examples. Most of the genes in the *mal* regulon, which encodes functions involved in maltose transport and utilization, were upregulated *in planta*. This regulon is typically encoded in Pantoea on LPP1, while in several pathogenic *Pantoea* species it is located on the chromosome (Fig. 3A).

**Figure 3.**
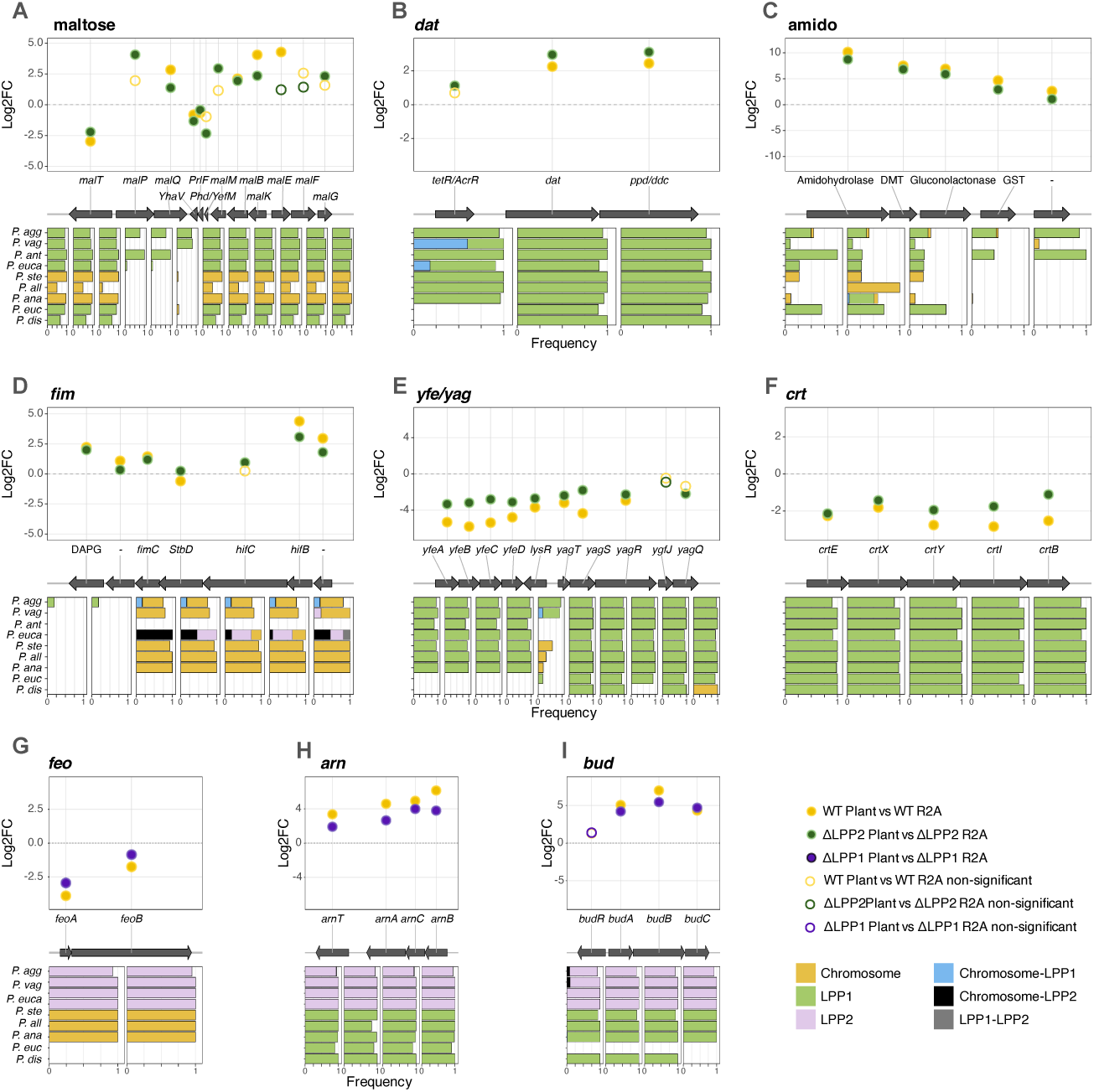
Representative plasmid loci exhibiting environment-dependent gene expression. (A–I) Representative examples of LPP1- and LPP2-encoded loci displaying differential expression between *in planta* and *in vitro* conditions. The upper panel shows the log2-fold change in gene expression *in planta* relative to R2A medium for the wild type, ΔLPP1, and ΔLPP2 strains. Each point represents a gene; filled symbols indicate significant differential expression as determined by DESeq2 (*P* < 0.05), whereas open symbols indicate genes that were not significantly differentially expressed. The corresponding genomic neighborhood is shown below, with arrows representing protein-coding genes. Stacked bar charts summarize the distribution of homologs among *Pantoea* species, with colors indicating the proportion of homologs encoded on the <u>chromosome, LPP1, LPP2, or multiple replicons in strains carrying more than one homolog.</u>

A locus containing diaminobutyrate aminotransferase (*dat*) and a pyridoxal phosphate-dependent decarboxylase (ppd/ddc) was upregulated *in planta* (Fig. 3B). Homologs of *dat* and *ppd/ddc* have been previously associated with symbiotic interaction in *Rhizobium leguminosarum* ^45^ and recently proposed to be involved in β-alanine biosynthesis in *Acinetobacter* ^46^. Homologs of both *dat* and *ppd/ddc* are universally encoded on LPP1 in *Pantoea* species (Fig. 3B). Another locus upregulated *in planta* encodes homologs of amidohydrolase and gluconolactonase (Fig. 3C). Members of these enzyme families have been implicated in the degradation of N-acyl homoserine lactones (AHLs) and are associated with quorum-sensing pathways or communication with eukaryotic hosts ^47^. In *P. agglomerans*, these genes occur on LPP1, whereas their homologs are chromosomally encoded in *P. stewartii* and *P. alii* (Fig. 3C). A locus encoding fimbrial proteins was overexpressed *in planta* (Fig. 3D). In *P. agglomerans* R1, homologous *fim/hif* genes are present on both LPP1 and the chromosome, with the chromosomal copy also showing increased expression *in planta*, albeit to a lesser extent (Table S3). The *fim/hif* locus is chromosomally encoded in most *Pantoea* species and can be found on LPP1 or LPP2 in several *P. agglomerans* and *P. anthophila* strains (Fig. 3D).

Examples of genes overexpressed under *in vitro* conditions include the *yfeABCD* (also termed *sitABCD*) iron uptake operon and the neighboring *yagTSRQ* (also termed *paoABCD*) operon encoding a xanthine dehydrogenase complex ^48^. Both loci are broadly conserved on LPP1 plasmids across the genus *Pantoea*, excluding *P. eucrina* and *P. dispersa* that are lacking *yfe* homologs (Fig. 3E). The *crt* carotenoid biosynthesis locus that is associated with *Pantoea*’s typical yellow hue ^49^ was downregulated *in planta* and is likewise widely conserved among LPP1 plasmids (Fig. 3E). The expression patterns observed for the wild type were similar to those observed in the ΔLPP2 strain.

A total of 82 LPP2 genes (56%) were differentially expressed *in planta* relative to *in vitro*. Several of the induced loci are associated with nutrient acquisition, cell envelope modification, and plant-associated metabolism. For example, the ferrous iron uptake operon *feoAB* was under-expressed *in planta* relative to *in vitro* conditions (Fig. 3G). Notably, the *feo* operon is located in LPP2 in all species harboring an LPP2 homolog, while in the other species it is located on the chromosome (Fig. 3G). Two loci comprising the *arn* operon (*arnTACB* and *arnFED*) were overexpressed *in planta* (Fig. 3H). Genes in *arn* operon are involved in lipopolysaccharide (LPS) modification and maintenance of the Gram-negative cell envelope ^50^, consistent with a role in membrane adaptation under host-associated conditions. Likewise, genes of the *bud* operon encoding enzymes for 2,3-butanediol biosynthesis were overexpressed *in planta*, indicating activation of a metabolic pathway previously linked to plant growth promoting bacteria ^51^. Indeed, LPP2 also encodes an *scrKYABR* operon for sucrose utilization (Table S3), further supporting a role of LPP2 in plant-associated carbohydrate metabolism. Both the *arn* and *bud* operons are located on LPP2 in the LPP2-hosts and on the LPP1 in the other species, with the exception of *P. dispersa* and *P. eucrina* where homologs of those operons were not found (Fig. 3I). The expression patterns of the above example operons were similar in the ΔLPP1 strain. Taken together, differentially expressed genes on LPP1 and LPP2 include multiple loci encoding traits that are often associated with an endophytic lifestyle.

### Plasmid curing alters co-resident plasmid gene expression depending on the environmental conditions

The highlighted examples indicate broad agreement in differential gene expression patterns between the wild type and strains lacking the co-resident plasmid. That is, most LPP1 genes that were differentially expressed *in planta* in the wild type were also differentially expressed in the ΔLPP2 strain. Nonetheless, several genes exhibited genotype-specific expression responses. For example, the LPP1 gene *malE* was upregulated *in planta* in the wild type but showed no significant differential expression between conditions in the ΔLPP2 strain (Fig. 3A). Approximately half of the LPP1 genes that were not differentially expressed *in planta* in the wild type exhibited differential expression *in planta* for the ΔLPP2 strain (Fig. S2). A similar pattern was observed for LPP2 genes, where approximately one third of the genes not differentially expressed *in planta* in the wild type were differentially expressed in the ΔLPP1 strain. These results indicate that plasmid curing alters the set of genes responding to environmental conditions.

The observed differences in gene expression are reflected in the overall plasmid gene expression profiles across genotypes. LPP1 gene expression was similar between the wild type and ΔLPP2 strains under *in vitro* conditions but diverged *in planta* (Fig. 4A). In contrast, LPP2 gene expression was similar between the wild type and ΔLPP1 strains *in planta* but differed under *in vitro* conditions (Fig. 4B). Statistical analysis supported the effects of both genotype and cultivation conditions on plasmid gene expression, with cultivation conditions as the primary determinant, affecting 356 (83%) LPP1 genes. Moreover, 231 (55%) LPP1 genes showed genotype-dependent expression differences, with 119 (28%) genes exhibiting genotype-specific responses depending on cultivation conditions (Fig. 4C). A similar pattern was observed for LPP2, where genotype affected expression of 64 (51%) LPP2 genes, with expression of 28 (22%) genes differing between the wild type and ΔLPP1 strains depending on the cultivation conditions (Fig. 4D). Taken together, these results reveal that plasmid curing alters gene expression of the co-resident plasmid in an environment-dependent manner.

**Figure 4.**
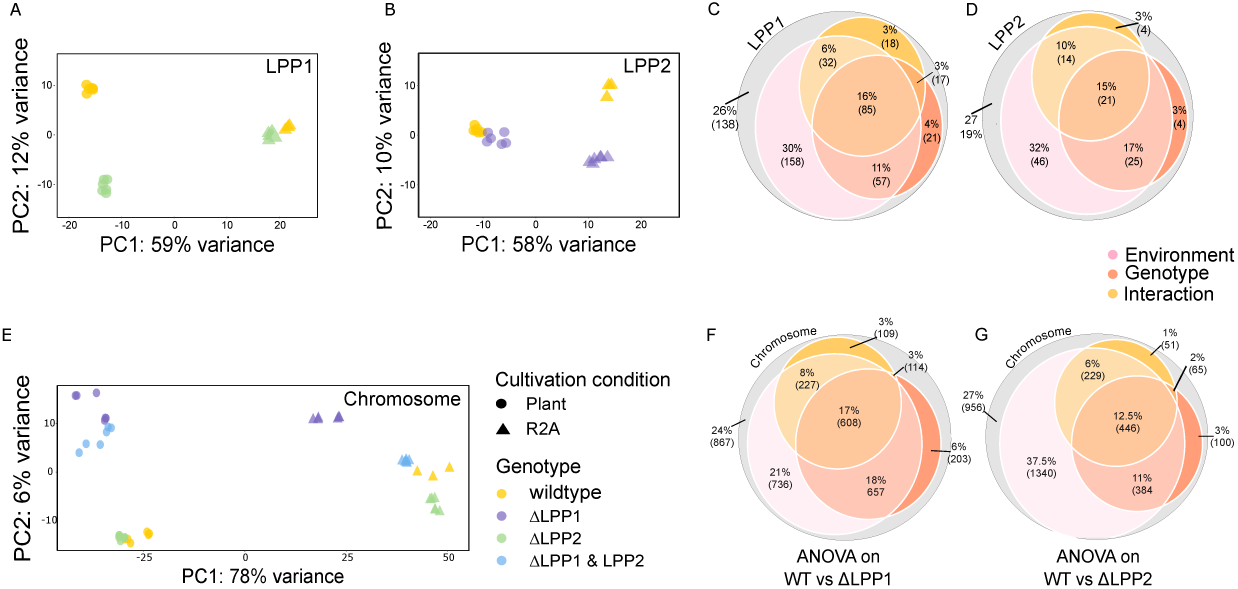
Plasmid curing alters plasmid and chromosomal gene expression in an environment-dependent manner. (A,B) Principal coordinates analysis (PCoA) of normalized transcript abundance for LPP1-encoded (A) and LPP2-encoded (B) genes across genotypes and growth conditions. Symbols represent independent biological replicates. **(C,D)** Summary of the effects of genotype and growth condition on LPP1 (C) and LPP2 (D) gene expression. Venn diagrams show the numbers of genes significantly affected by genotype, growth condition, or their interaction based on two-way ANOVA. **(E)** Principal coordinates analysis of normalized transcript abundance for chromosomal genes across genotypes and growth conditions. **(F,G)** Effects of plasmid curing and growth condition on chromosomal gene expression, comparing ΔLPP1 with the wild type (F) and ΔLPP2 with the wild type (G). Differentially expressed genes were identified by two-way ANOVA <u>(complete ANOVA results are provided in Table S3).</u>

### LPP1 curing leads to wide-range alteration of chromosomal gene expression *in planta*

Differences in gene expression depending on the genotype were observed also for chromosomal genes. The majority (65-70%) of chromosomal genes differentially expressed *in planta* in the wild type were also differentially expressed in the plasmid-cured strains. In addition, approximately 40% of chromosomal genes that were not differentially expressed in the wild type were differentially expressed in the plasmid-cured genotypes (Fig. S2). The effect of plasmid curing on chromosomal gene expression varied across genotypes and cultivation conditions (Fig. 4E). Under *in vitro* conditions, chromosomal gene expression profiles were similar between the wild type, ΔLPP2, and ΔLPP1&2 strains, but differed slightly from those of the ΔLPP1 strain. *In planta*, chromosomal gene expression profiles of the ΔLPP1 and ΔLPP1&2 strains clustered together and were distinct from the wild type and ΔLPP2 strains. Together, these results indicate that plasmid curing alters chromosomal gene expression in an environment-dependent manner.

Further statistical analysis reveals that cultivation conditions are the primary determinant of chromosomal gene expression, affecting most (ca. 85%) chromosomal genes. In addition, expression of approximately 60% of chromosomal genes differed significantly between the wild type and the ΔLPP1 strain, whereas 40% differed between the wild type and the ΔLPP2 strain (Fig. 4FG). These results indicate a stronger effect of LPP1 curing on chromosomal gene expression in planta, consistent with the larger negative fitness effect of LPP1 curing on Pantoea growth *in planta* (Fig. 1E).

Curing of LPP1 has previously been shown to reduce motility in *P. vagans* ^32^. In line with this observation, *flgK* and *flgL* of the *flg* regulon, which are upregulated *in planta* in the wild type, were downregulated in the ΔLPP1 strain (Fig. 5A). Likewise, all genes of the *che* operon that were overexpressed *in planta* in the wild type were downregulated in the ΔLPP1 strain (Fig. 5B). A genotype-specific effect on *flg* and *che* gene expression was also observed, albeit more weakly, in comparisons between the wild type and ΔLPP2 strains (Table S3). In line with the differential expression level, all plasmid-cured *Pantoea* strains exhibited reduced motility relative to the wild type (Fig. S3).

**Figure 5.**
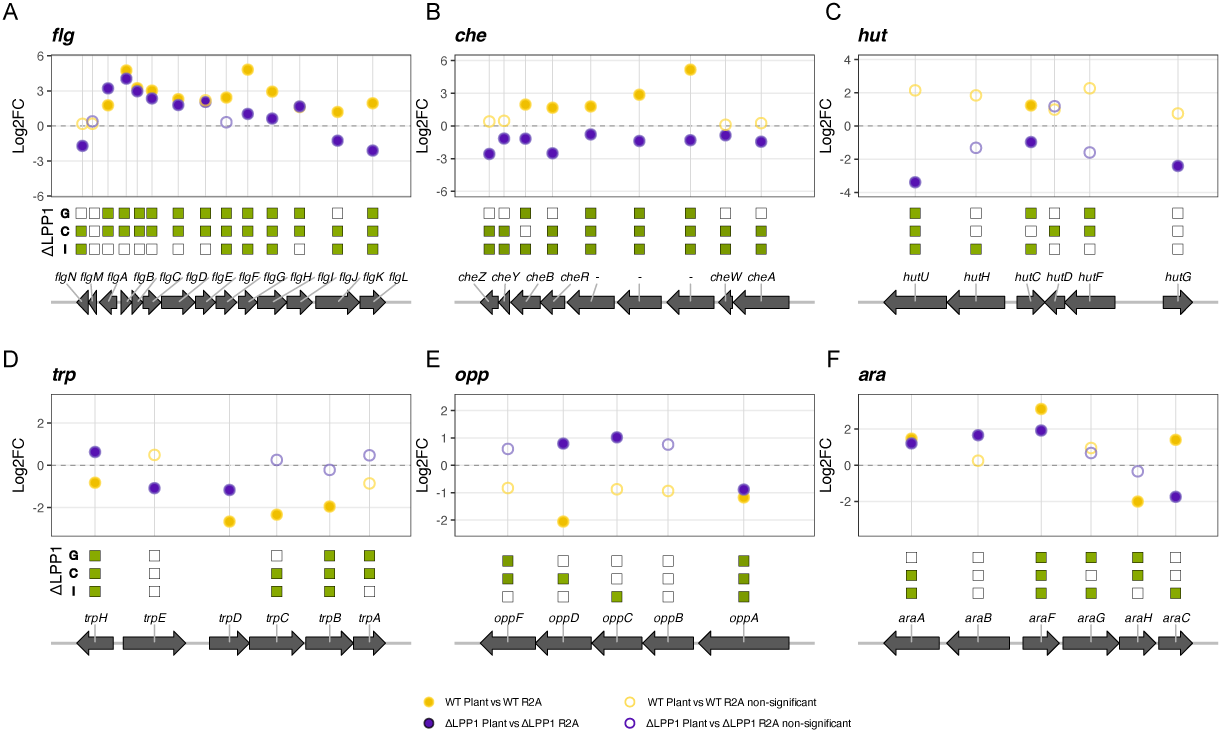
Representative chromosomal loci exhibiting LPP1-dependent, environment-specific gene expression. (A–F) Representative chromosomal loci displaying differential expression between *in planta* and *in vitro*conditions in the wild type and ΔLPP1. The upper panel shows the log-fold change in gene expression *in planta* relative to R2A medium. Each point represents a gene; filled symbols indicate significant differential expression as determined by DESeq2 (*P* < 0.05), whereas open symbols indicate genes that were not significantly differentially expressed. Boxes below each plot summarize the results of the two-way ANOVA, with green boxes indicating a significant effect of genotype (G), growth condition (C), or their interaction (I). The corresponding genomic neighborhood is shown beneath each plot, with arrows representing protein-coding genes. All presented loci are core chromosomal genes with only few exceptions.

Several chromosomally encoded metabolic genes were differentially expressed between the wild type and ΔLPP1 depending on the cultivation conditions. Examples include the histidine utilization (*hut*) operon, which functions in L-histidine catabolism. In particular, expression of the *hut* operon repressor, *hutC*, was lower *in planta* for ΔLPP1 compared to the wild type (Fig. 5C). Differences in expression level were also observed for genes of the tryptophan biosynthesis (*trp*) operon, most of which were more highly expressed *in vitro* than *in planta* (Fig. 5D). In addition, several genes of the oligopeptide permease (*opp*) operon, which mediates oligopeptide uptake, were overexpressed *in planta* in the ΔLPP1 strain but not in the wild type (Fig. 5E). Differential expression between the wild type and ΔLPP1 was also observed for genes of the L-arabinose utilization (*ara*) operon. Notably, the operon’s main regulator, *araC*, was overexpressed *in planta* in the wild type, whereas in the ΔLPP1 strain it was overexpressed under *in vitro* conditions (Fig. 5F). Together, these examples indicate that LPP1 curing affects the expression of chromosomally encoded genes involved in transport and metabolism of amino acids and carbohydrates.

### Plasmid curing reduces the capacity to utilize diverse carbon sources

To further investigate the effects of plasmid curing on the metabolic capacity of Pantoea, we compared substrate-dependent respiration of the wild type and plasmid-cured strains across individual carbon sources using OmniLog© phenotype microarrays. A total of 95 carbon sources were tested, of which 56 elicited detectable respiration in the wild type. Across these substrates, the wild type generally exhibited a higher inferred maximum respiration rate than the plasmid-cured strains. In contrast, the ΔLPP1 strain showed no detectable respiration on 32% of the substrates metabolized by the wild type (Fig. 6A). For example, ΔLPP1 showed no detectable respiration on maltose (Fig. 6B), consistent with the presence of the *mal* regulon on LPP1 and previous reports ^28,32^. By contrast, ΔLPP1 and the wild type exhibited similar respiration dynamics when supplied with arabinose (Fig. 6C), in agreement with the chromosomal location of the *ara* operon (Fig. 5F). However, ΔLPP1 displayed a pronounced delay in respiration relative to the wild type and ΔLPP2 when supplied with succinic acid (Fig. 6E). Respiration of ΔLPP1 was likewise delayed or undetectable for additional substrates, including L-glutamine, L-glutamic acid, fumaric acid, L-lactic acid, L-malic acid, D-malic acid, or L-alanine was supplied as the sole carbon source (Fig. 6A,D,F). Notably, these phenotypes involve substrates whose utilization is not exclusively dependent on LPP1-encoded pathways.

**Figure 6.**
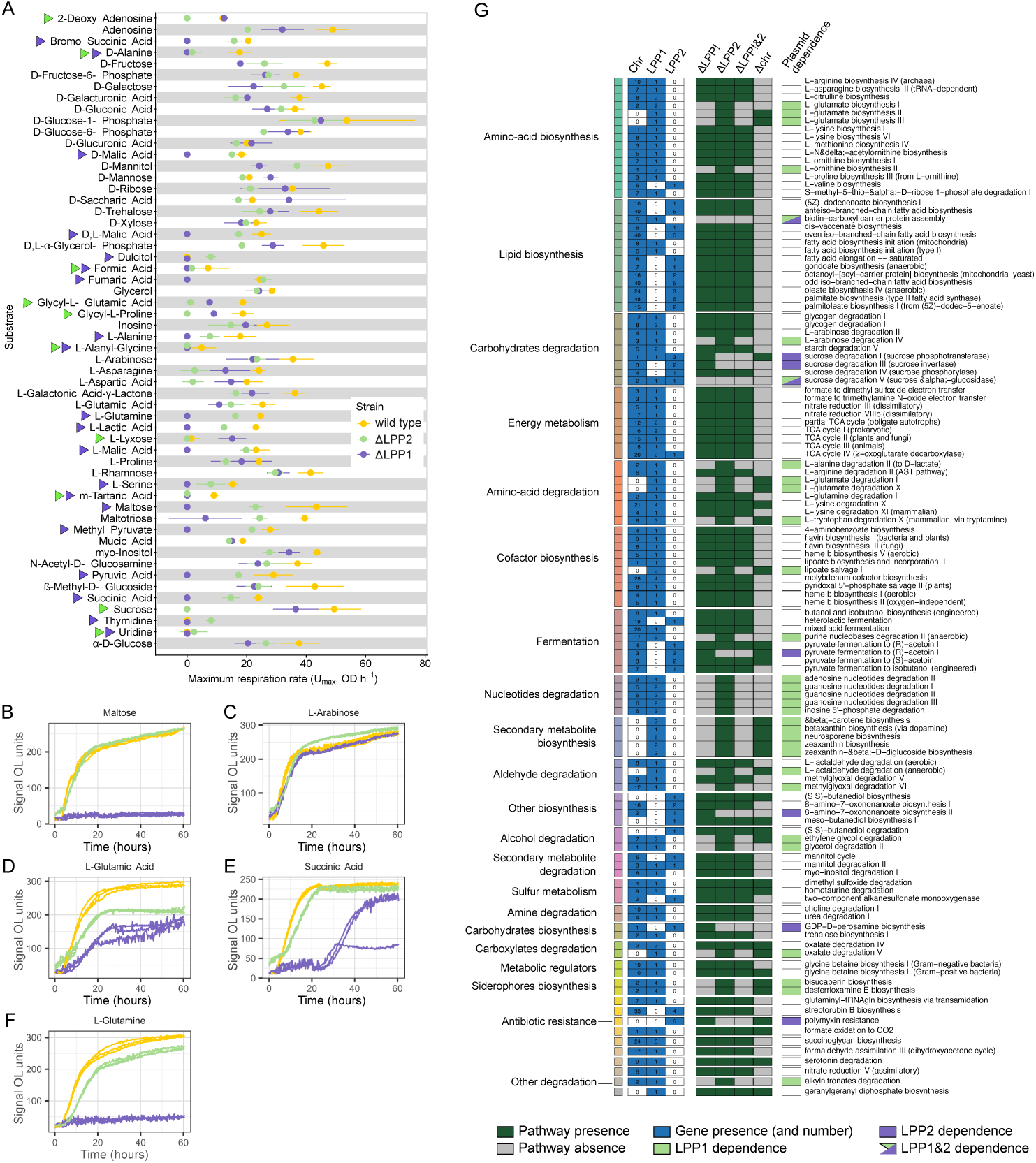
Plasmid curing alters carbon utilization and reveals plasmid-dependent metabolic pathways. **(A)** Maximum substrate-dependent respiration rate (*r_max_*) of wild-type *Pantoea agglomerans* R1 and plasmid-cured derivatives across individual carbon sources, measured using Biolog phenotype microarrays. **(B–F)** Representative respiration kinetics of the wild type and plasmid-cured strains on selected carbon substrates. **(G)** Reconstruction of genome-scale metabolic models for the wild type and plasmid-cured strains. Metabolic pathways containing plasmid-encoded genes are grouped by functional category (left). The left panel shows the distribution of pathway-associated genes among the chromosome, LPP1, and LPP2, with numbers indicating the number of reactions encoded on each replicon. The middle panel summarizes the presence or absence of each pathway in metabolic models reconstructed for the wild type, ΔLPP1, ΔLPP2, ΔLPP1&2, and a chromosome-free reference model containing only plasmid genes (ΔChromosome). Green boxes indicate the presence of a pathway in the corresponding model. The right panel classifies pathways according to plasmid dependency: pathways present in the wild-type model but absent from both the ΔLPP1 and ΔLPP1&2 models were classified as LPP1-dependent, whereas pathways present in the wild-type model but absent from both the ΔLPP2 and ΔLPP1&2 models were classified as LPP2-dependent.

The ΔLPP2 strain exhibited respiration dynamics broadly similar to those of the wild type, with only a few exceptions. No detectable respiration was observed when ΔLPP2 was supplied with sucrose (Fig. 6A), consistent with the presence of the *scr* operon on LPP2 and previous reports ^32^. In addition, ΔLPP2 displayed delayed respiration when supplied with D-alanine or L-alanine (Fig. 6A). Because genes for alanine catabolism (*dadAX*) are chromosomally encoded, the reduced ΔLPP2 respiration further supports that plasmid curing may affect chromosomally encoded metabolic functions. Respiration of ΔLPP2 in response to individual nitrogen sources was likewise broadly similar to that of the wild type (Fig. S4). The effects of nitrogen sources on ΔLPP1 metabolism could not be assessed because the assay relies on succinate as the sole carbon source, which elicits no detectable respiration in this strain. Nonetheless, the assay revealed that D-glucosamine and adenosine can substitute for succinate as carbon sources and restore detectable respiration in ΔLPP1 (Fig. S4).

Taken together, plasmid curing reduces the capacity of Pantoea to utilize diverse carbon sources, with LPP1 curing having the stronger effect. Moreover, impaired respiration on substrates whose utilization does not directly depend on plasmid-encoded pathways suggests metabolic or regulatory interactions between the plasmids and chromosome, consistent with the observed effects of plasmid curing on chromosomal gene expression.

### Genome-scale metabolic modeling identifies plasmid-dependent metabolic pathways

To gain further insight into the contribution of LPP1 and LPP2 to the metabolic capacity of Pantoea, we reconstructed a genome-scale metabolic model of Pantoea and examined the distribution of plasmid-encoded genes across metabolic pathways. LPP1 genes were predicted to participate in numerous pathways, primarily those involved in amino acid metabolism, carbohydrate degradation, energy metabolism, cofactor and secondary metabolite biosynthesis, as well as nucleotide and aldehyde degradation. LPP2 genes were associated with a smaller number of pathways, including lipid biosynthesis, carbohydrate degradation, and fermentation (Fig. 6G). Most pathways containing plasmid-encoded genes also included chromosomally encoded functions. However, 13 pathways were supported exclusively by LPP1-encoded genes. Examples include L-glutamate biosynthesis II and III, L-glutamate degradation I and X, and five secondary metabolite biosynthesis pathways, including carotenoid biosynthesis. In contrast, four pathways comprised exclusively LPP2-encoded genes, including pathways associated with polymyxin resistance, (2S,3S)-butanediol biosynthesis and degradation, and meso-butanediol I biosynthesis. Together, these results suggest that LPP1 and LPP2 contribute to a broad range of metabolic functions.

To identify metabolic pathways that depend on plasmid-encoded functions, we reconstructed metabolic models of the plasmid-cured strains by excluding plasmid genes from the corresponding models. In addition, we generated a hypothetical Δchromosome model in which chromosomal genes were excluded (Fig. 6G). Comparison of the four models confirmed that most pathways composed exclusively of plasmid-encoded genes depend on the presence of the corresponding plasmid. For example, L-glutamate biosynthesis and degradation, carotenoid biosynthesis, and lipoate salvage were predicted to depend on LPP1 genes. An exception was the geranylgeranyl diphosphate biosynthesis pathway, which contains a single LPP1-encoded gene but was predicted to remain functional in all models. Only a few metabolic pathways were predicted to depend on LPP2 genes, including sucrose degradation, polymyxin resistance, pyruvate fermentation, 8-amino-7-oxononanoate biosynthesis, and GDP-D-perosamine biosynthesis. Notably, all lipid biosynthesis pathways containing LPP2 genes, with the exception of biotin biosynthesis, were predicted to remain functional in the absence of LPP2. These results indicate that, despite contributing genes to several lipid biosynthesis pathways, LPP2 is predicted as generally dispensable for their function.

Most pathways containing both plasmid- and chromosome-encoded genes were retained in the models of the plasmid-cured strains (e.g., lysine and valine biosynthesis pathways; Fig. 6G). This pattern suggests that essential reactions in these pathways are encoded on the chromosome, whereas plasmid genes either perform redundant functions or contribute to accessory reactions. In contrast, pathways containing both plasmid- and chromosome-encoded genes that were absent from the cured-strain models likely depend on plasmid genes for essential pathway functions. Examples include L-glutamate biosynthesis I, L-ornithine biosynthesis II, L-arabinose degradation, sucrose degradation II and III, L-alanine degradation II, and glycerol degradation II. Notably, two pathways comprising both plasmid and chromosomal genes were predicted to be absent from all four *in silico* cured models: sucrose degradation V and biotin-carboxyl carrier protein assembly biosynthesis. These pathways may depend on essential transporters encoded by plasmid genes. Further examination of plasmid-genes in LPP1-dependent pathways identified glutamate dehydrogenase (*gdhA*) as a common component of multiple pathways (Table S4). This enzyme catalyzes reactions in the L-glutamate biosynthesis I–III, L-glutamate degradation I, L-alanine degradation II (to D-lactate), and L-ornithine biosynthesis II pathways. Among these pathways, *gdhA* emerged as a recurrent component, linking several LPP1-dependent functions involved in amino acid metabolism.

## Discussion

Plasmids encoding traits that promote association with eukaryotic hosts represent a distinctive class of plasmids whose evolutionary dynamics differ from those of transient accessory plasmids. Host-associated environments impose persistent selection for the maintenance of plasmids supplying symbiosis traits, in contrast to the episodic selection acting on accessory plasmids. Long-term plasmid retention under continuous selection is expected to increase opportunities for regulatory and metabolic integration with the chromosome, ultimately promoting the evolution of chromosome-like properties and plasmid domestication ^9,10^.

Several LPP1 loci that are upregulated *in planta* are likely to encode functions that contribute to plant association, and most are widely conserved in LPP1 across the genus. Among these, the LPP1-encoded *mal* regulon is a strong candidate for promoting the utilization of plant-derived carbohydrates. Maltose is present in root exudates ^52^ and can be generated through the degradation of plant starch, providing a potential carbon source in the rhizosphere ^53^. Moreover, maltose and maltodextrin utilization are defining traits of heritable seed-borne bacteria in wheat^54^. Likewise, the environmentally induced *yfe/yag* locus suggests a role for LPP1 in iron acquisition during host association. Competition for iron is a well-established determinant of bacterial fitness in plants and is a characteristic feature of plant-associated bacteria ^55^ and the wheat seed microbiome ^54^. Because the *in planta* transcriptomes were contrasted with those of R2A-grown cultures, some of these expression differences may reflect changes in nutrient availability, oxygenation, or growth rate rather than responses to plant-specific cues. Such environmental differences represent conditions encountered during plant colonization, and the functions encoded by these responsive LPP1 loci are consistent with traits expected to enhance bacterial fitness in the plant environment. Together, these examples suggest that LPP1 promotes *Pantoea* association with plants through a diverse repertoire of metabolic and colonization-related functions rather than through a single adaptive trait.

Although curing LPP2 had no detectable effect on *Pantoea* colonization of wheat seedlings, the *in planta* expression pattern of LPP2 genes highlights several loci encoding functions widely linked with plant-associated bacteria. These include the ferrous iron uptake operon (*feo*), which contributes to host colonization in both plant pathogens ^56^ and symbionts such as *Bradyrhizobium japonicum* ^57^. Likewise, the LPP2-encoded *scrKYABR* sucrose utilization operon is reminiscent of sucrose catabolism loci described in phytopathogens such as *Erwinia amylovora* ^58^. Notably, these functions are commonly chromosomally encoded in plant-associated bacteria, whereas in *Pantoea* R1 they occur in close linkage on LPP2. Thus, similar to LPP1, LPP2 carries a repertoire of genes associated with host-associated lifestyles. The absence of a detectable colonization phenotype following LPP2 curing therefore suggests that these functions are either dispensable under the conditions examined or contribute to aspects of host association not captured by our colonization assay.

The transcriptional consequences of plasmid curing further indicate that the contribution of LPP1 and LPP2 to *Pantoea* physiology extends well beyond the functions encoded by plasmid genes. Curing either plasmid significantly altered chromosomal gene expression, and these effects depended on the environmental conditions. These observations demonstrate regulatory cross-talk between the domesticated plasmids and the chromosome. Similar genome-wide transcriptional responses have been reported following the curing of stably inherited plasmids, including the virulence plasmids of *Rhodococcus equi* and *Shigella flexneri* and the secondary replicons of *Dinoroseobacter shibae* and *Sinorhizobium meliloti* ^59–62^. More generally, acquisition of plasmids is often accompanied by changes in chromosomal gene expression, including metabolic pathways ^63–65^. In newly acquired plasmids, such transcriptional perturbations are commonly interpreted as regulatory conflicts that contribute to the fitness costs of plasmid carriage ^66,67^. In contrast, LPP1 and LPP2 have been vertically inherited over evolutionary timescales and exhibit multiple hallmarks of long-term coevolution with their host ^29^. The regulatory interactions observed here reflect functional integration of LPP1 and LPP2 into the transcriptional architecture of *Pantoea*.

The reduced metabolic capacity observed in the plasmid-cured strains mirrors the changes in transcriptional profiles, indicating that regulatory integration is translated into genome-wide metabolic integration. Curing LPP1 substantially reduced the ability of *Pantoea* to utilize diverse carbon sources. Similar metabolic effects have been reported following plasmid curing in rhizobia, including *Rhizobium leguminosarum* ^68^. Genome-scale metabolic reconstruction suggests that dependence on plasmid-encoded functions arises through two complementary mechanisms. First, a small number of pathways depend directly on plasmid-encoded genes, including maltose utilization and several amino acid biosynthetic and degradation pathways. Second, and more importantly, most metabolic pathways containing plasmid genes also comprise chromosomally encoded enzymes, indicating that the plasmid and chromosome jointly encode mosaic metabolic networks rather than independent modules. Integration of plasmid- and chromosome-encoded genes into a common metabolic pathway has so far been described in only a few cases (e.g., *Pseudomonas* ^69,70^). Among the plasmid-encoded enzymes, glutamate dehydrogenase (*gdhA*) emerges as a central metabolic hub linking glutamate and ornithine metabolism with carbon flux into the tricarboxylic acid cycle ^71–73^. We hypothesize that this organization provides a mechanistic explanation for the disproportionate metabolic consequences of LPP1 curing, despite the relatively small number of plasmid-specific pathways. Indeed, several growth deficiencies observed experimentally were not predicted by the metabolic model, suggesting that the metabolic effects of plasmid curing extend beyond the direct loss of enzymatic reactions. These discrepancies are likely explained by the regulatory interactions between the plasmid and chromosome revealed by the transcriptomic analyses. Such interactions may be mediated by plasmid-encoded DNA-binding proteins, including H-NS-like proteins or broad-acting transcriptional regulators such as members of the LysR family ^74,75^. Together, our findings indicate that the contribution of LPP1 to host fitness arises primarily from its metabolic integration with the chromosome rather than from the simple acquisition of accessory metabolic genes.

Our findings identify LPP1 as a clear example of a chromosome-like symbiosis plasmid. LPP2 likely represents a similar evolutionary trajectory, although the ecological conditions under which it confers a fitness advantage remain to be determined. Although both plasmids remain genetically separable from the chromosome and can be experimentally cured under laboratory conditions, they exhibit multiple hallmarks of plasmid domestication, including stable vertical inheritance ^29^, extensive regulatory and metabolic integration with the chromosome, and in the case of LPP1, a substantial contribution to host fitness in its natural environment. These observations suggest that chromosome-like plasmids, or chromids, should not be defined solely by their dispensability under laboratory conditions. Rather, they should be viewed as replicons that have acquired chromosome-like properties through long-term coevolution with their host. From this perspective, plasmid domestication represents a continuum of evolutionary integration, with LPP1 exemplifying an advanced stage in the transition from a mobile genetic element to a chromosome-like replicon that has become an integral component of the bacterial genome.

## Methods

### Generation of plasmid cured strains

The LPP1 was cured by thermal destabilization as described by Smits et al (2011). Briefly, a culture of the wild-type strain was incubated at 37°C in LB medium overnight and 10^-5^ dilutions were plated onto LB agar and incubated at 37°C for 36 h. As LPP1 carries the carotenoid biosynthesis cluster *crtZEXYIB* (GQQ16_018785-GQQ16_018810), colonies that had lost the plasmid were identified by their non-pigmented phenotype. Loss of LPP1 was verified by PCR using LPP1-specific primers (repB_1-fw-R1-LPP1 and repB_1-rv-R1-LPP1; Table S5), with genomic DNA from the wild-type strain serving as a positive control. Two candidate cured clones lacking the LPP1-specific amplicon were subsequently confirmed by whole-genome sequencing. Curing of LPP2 was achieved by substituting the *arn*-locus (*arnBCADTEF*, GQQ16_021980–GQQ16_021950) on LPP2 with a [*sacB nptII*] cassette which confers sucrose sensitivity and kanamycin resistance. For this purpose, plasmid pGT43-arn-sub, harboring the cassette, was introduced into wild-type *P. agglomerans* strain R1 and its ΔLPP1 derivative using *E. coli* MFD*pir* as the donor. Conjugations were set up by mixing 500 µl of overnight cultures of *E. coli* MFD*pir* pGT43-*arn*-sub with 500 µl of each *Pantoea* strain. Fifty-microliter drops of mixed cultures were spotted onto non-selective LB agar supplemented with 0.1 mM diaminopimelic acid (DAP) to enable growth of *E. coli* MFD*pir*. The plates were incubated for 6 h at 30 °C. After incubation, the agar patches were excised and resuspended in 1 ml PBS. The recovered bacteria were plated on LB agar supplemented with 10 µg/ml kanamycin lacking DAP to select for *Pantoea* transconjugants and against *E. coli* MFD*pir*, and incubated at 30°C overnight. Resulting colonies were streaked on LB agar supplemented with 10 µg/ml kanamycin and, in parallel, on LB agar supplemented with 10 µg/ml chloramphenicol to screen for co-integration of the plasmid backbone. Colonies resistant to kanamycin and sensitive to chloramphenicol were screened by colony PCR (primers are listed in Table S5).

To cure LPP2 from *Pantoea,* 25 µl overnight cultures of the wild type and the ΔLPP1 strain both carrying the Δ*arnBCADTEF*::[*sacB nptII*] allele were plated on R2A agar plates supplemented with 5 % sucrose and incubated at 30°C overnight to select for loss of LPP2. Resulting colonies were streaked on non-selective R2A agar and, in parallel, on R2A agar supplemented with 10 µg/ml kanamycin. Clones resistant to sucrose and susceptible to kanamycin were screened by colony PCR targeting the *repB* gene of LPP2 (primers are listed in Table S5). Finally, loss of LPP2 was verified by whole genome sequencing.

### Detection of chromosomal variants in plasmid cured strains

The presence of chromosomal variants in the plasmid-cured strains was tested by whole genome sequencing. For sequencing single colonies of wild-type and plasmid-cured *P. agglomerans* R1 strains were picked to inoculate overnight cultures. From the cultures, DNA was extracted using the WIZARD DNA extraction kit (Promega) as instructed by the manufacturer. The acquired DNA was sequenced on Illumina MiSeq 2500 (Eurofins Scientific SE, Lux) with 2X150 bp paired end reads.

Paired-end reads were trimmed by quality using trimmomatic v0.39 ^76^. Adapter sequences were removed using the provided adapter file (ILLUMINACLIP:2:30:10). Low-quality regions were trimmed using a 4-bp sliding window with a minimum average Phred quality score of 20 (SLIDINGWINDOW:4:20), and reads shorter than 50 bp after trimming were discarded (MINLEN:50). Variants were first identified for the wild-type and cured strains by comparison with the reference genome of *Pantoea agglomerans* R1 (NCBI accession GCF_010667995.1) using breseq v0.39.0 in polymorphism mode ^77^. To prevent spurious read alignments to plasmid sequences that had been experimentally cured, plasmid contigs have been excluded from the reference genome. Only variants with an allele frequency of at least 5% were retained. No nonsynonymous mutations were shared between ΔLPP1 and ΔLPP1&2, except for a fixed 9-bp tandem duplication in the envelope stress sensor histidine kinase gene *cpxA*. Previous studies have shown that deletion of *cpxA* has no detectable fitness effect during *in vitro* growth in *Escherichia coli* or *Vibrio alginolyticus* ^78,79^. Thus, the *cpxA* variant is unlikely to account for the reduced fitness of the ΔLPP1 and ΔLPP1&2 strains *in vitro*. No nonsynonymous mutations unique to ΔLPP1 were detected. Nonsynonymous mutations unique to ΔLPP1&2 were infrequent and generally present at low allele frequencies (<10%), with the exception of a fixed mutation in a gene encoding an efflux transporter outer membrane subunit. The complete set of detected variants is provided in Supplementary Table S1.

### Bleaching and maintenance of germ-free wheat plants in colonization experiments

A total of 50 grains of the wheat cultivar Benchmark were submerged in 50 ml 12% sodium hypochlorite with 0.05 % sterile filtered Tween 20 for 30 minutes with gentle agitation and subsequently washed three times with sterile water (as previously described ^35^). The grains were individually placed in 24-well plates with plant media (modified *Arabidopsis/P. indica* co-cultivation medium ^80^) supplemented with 15 µg/ml neomycin, 25 µg/ml streptomycin, 25 µg/ml spectinomycin and 6 µg/ml amphotericin B and incubated in the dark at 25°C. After three days, germinated grains were transferred individually into 6-well-plates with 5 ml plant media and incubated in a growth chamber for an additional two to three days. The media was replaced daily throughout all incubation steps. Once the roots had reached a length of approximately 1.5 cm, the sterile seedlings were used for colonization experiments.

### Single colonizer experiments

Inoculation media was prepared by diluting fresh over-night Pantoea cultures carrying pRL153-GFP in plant medium to the desired density. Dilutions were prepared based on the optical density of the over-night cultures to a known standard (i.e., OD of representative overnight cultures of Pantoea with known viable titer). For each experiment the exact inoculum density was confirmed by plating on R2A media plates.

The sterile seedlings were placed onto plastic cell strainers with a 5 mm hole and individually placed in 6-well-plates with 5 ml of inoculated plant media containing 200 CFU / ml (i.e., 2000 cells per plant). Five hours after inoculation and then every 24 h, the plants were transferred to new 50 ml tubes with plant medium supplemented with 10 µ/ml neomycin and 6 µg/ml amphotericin B, and six randomly chosen plants were sampled. The inoculated plants were transferred into a growth chamber (MLR-352H, Panasonic) and were incubated for 24 hours under following conditions: 16 h/8 h (light/dark) at 25°C/20°C with 65% relative humidity. For sampling roots were placed on sterile filter paper to absorb excess plant media and then separated at the junction to grain/seed using a sterile scalpel. The roots were placed individually in 1.5 ml microcentrifuge tubes, and their sample mass was recorded before manual homogenization with a sterile pestle. One milliliter of sterile PBS containing 0.05% Tween 20 was added to each sample, followed by vigorous vortexing for 1 minute. Samples were serially diluted, plated on non-selective R2A plates, and incubated at 25°C overnight. Fluorescent colonies were counted to estimate Pantoea population size recovered from the root system. Experiments were continued for 120h post inoculation.

### In planta competition assays

For competitions assays, wild-type Pantoea tagged with pTW1-mcherry and plasmid cured Pantoea strains tagged with pRL153-GFP were prepared as described above. Plant medium was inoculated with the competing strains at a 1:1 ratio to a final density of 400 CFU/ml (200 CFU/ml per strain). Seed sterilization, inoculation and sampling was carried out as described for single colonization experiments.

### Fitness evaluation via competitions *in vitro*

Fitness competitions between wild-type Pantoea and the plasmid-cured strains ΔLPP1, ΔLPP2, and ΔLPP1&2 were performed in R2A broth with four biological replicates. The wild type was tagged with pTW1-mCherry, whereas the plasmid-cured strains were tagged with pRL153-GFP. Previous experiments using this system showed that the fitness effects of both reporter plasmids are negligible, although pTW1-mCherry imposed a slightly higher fitness cost than pRL153-GFP ^35^. Because the wild type carried pTW1-mCherry, any resulting bias would favor the plasmid-cured strains; thus, estimates of the fitness advantage of the wild type are expected to be conservative.

For each experiment, overnight cultures grown from a single colony of the respective strain were diluted 1:100 in R2A medium and mixed in a 1:1 volumetric ratio. Aliquots of 0.5 ml of the mixture were transferred to 2-ml deep-well plate, corresponding to an initial density of approximately 2 × 10^6^ cells/ml for each competitor. Mixed cultures were incubated overnight at 30°C with shaking and subsequently transferred daily by 1:100 dilution into fresh R2A medium for five consecutive days under the same incubation conditions. At the onset of the experiment and after each transfer, the relative abundance of the two competing strains was determined by non-selective plating. Plasmid-carrying colonies were identified by their emitted fluorescence signal upon excitation with a green-blue LED flashlight (Nippon Genetics Europe) while wild-type colonies remained non-fluorescent. The selection coefficient *s*, which is equal to the selective difference between the two competing strains during each transfer, was calculated as *s* = *b*/ln(1/*d*) as previously described ^81^, where *b* is the slope of ln(plasmid-cured strain/wild type) as a function of transfer, and *d* is the dilution factor (1:100). Wild-type fitness was set to 1.0, and the value of *w* (relative Darwinian fitness) for the plasmid-carrying strain was calculated as *w* = 1 + *s*.

### RNA extraction and sequencing

Germ-free wheat plants were grown as previously described in *Soluch et. al*. Fifty millliters of a culture of the respective *P. agglomerans* strain were added to 1L of plant medium and germinated seedlings were then transferred to this inoculated plant medium as described above. The inoculated plants were transferred into a growth chamber (MLR-352H, Panasonic) and were incubated for 24 hours under the following conditions: 16 h/8 h (light/dark) at 25°C/20°C with 75% relative humidity. Every 24 h, the plants were transferred to new tubes containing fresh plant medium. After 48 h post-inoculation, roots from 30-40 plants were collected in 1.5 ml microcentrifuge tubes containing ice-cold saline with Tween 20. Bacteria were recovered from the root surface by shaking in a Precellys mixer at 4000 rpm for 30 s and collecting the supernatant. Plant debris was removed by centrifugation at 1.1 × g for 1 min at 4°C, followed by collection of the supernatant. Finally, bacterial cells were harvested by centrifugation of this supernatant at 9.000 × g for 1 minute. Cells were then resuspended in saline containing Tween 20. For *in vitro* samples, all genotypes were grown in R2A medium at 25°C with shaking at 200 rpm. Cells were harvested during mid-exponential growth (OD600-0.3-0.4) by centrifugation at 9000 × g in a cold centrifuge and resuspended in ice-cold saline with Tween 20. Two biological replicates, each comprising three technical replicates, were included for each sample.

RNA extraction from both *in planta* and *in vitro* bacterial samples was performed according to the Direct-zol RNA MiniPrep protocol (Zymo Research). The RNA preparations were additionally treated with RNase-free DNase I to remove residual DNA and subsequently purified using the RNA Clean & Concentrator-25 kit (Zymo Research). RNA was eluted in RNase free-water and stored at −80°C until further use. All steps were performed on ice unless stated otherwise to minimize RNA degradation. RNA concentration and quality were assessed using a Nanodrop spectrophotometer (ThermoFisher Scientific), a Qubit fluorometer with RNA High Sensitivity reagents (ThermoFisher Scientific), and an Agilent Bioanalyzer. High-quality RNA samples were then selected for sequencing. mRNA enrichment and plant RNA depletion were performed using the Ribo-Off kit (Vazyme Biotech). For RNA sequencing, PE150 libraries were prepared with the Trio RNA-Seq library kit (NuGEN Technologies), and sequenced by BGI Genomics Co., Ltd. on a DNBSEQ platform, generating up to 3 Gb of clean sequence data per sample.

### RNA sequences assembly and analysis

The sequencing reads were demultiplexed by the sequencing facility. The reads were trimmed with Trim Galore (parameters – paired, quality-30, length-20, max_n-5). The quality of the reads was checked using FASTQC (http://www.bioinformatics.babraham.ac.uk/projects/fastqc/). Reads were mapped onto the genomes of *P. agglomerans* R1 (Project accession <u>WSSS00000000.1</u>) using Bowtie (bowtie2-2.3.2) and Samtools. Transcript level abundance for each sample was estimated using salmon. The quants obtained from salmon were made accessible for downstream analysis using Tximport. Differential gene expression analysis was carried out using DESeq2. Transcripts per million (TPM) were obtained for each sample from the salmon output files, normalized between and across samples and log transformed for further analysis. Raw TPM values obtained from DEseq2 are supplied in Table S6. A two-way ANOVA was performed on the log transformed TPMs of each gene individually to evaluate and compare the effect of genotype versus condition on gene expression (p<5%), followed by correction for multiple comparisons using FDR. Finally, COG annotations were obtained using EggNOG mapper and the enrichment analysis for COG categories was performed using Fisher’s exact test (p<5%) followed by an FDR correction.

### Pangenome analysis

The pangenome analysis of the *Pantoea* genus was done using the gene families previously published ^29^. Here, we studied the frequency of genes in the subset of 201 genomes carrying LPP1 and LPP2 -the later, only in species where the LPP2 is expected to be present. All assemblies used in the analysis have a completeness ≥ 97% except for one *P. ananatis* assembly with a completeness of 86%. The pangenome data is supplied in Table S7.

### Carbon and Nitrogen utilization assay

All four genotypes (WT, ΔLPP1, ΔLPP2, ΔLPP1&2) were grown overnight in R2A medium at 30°C and shaking at 200 rpm. The cells were washed in PBS 2 times and resuspended in IF-0 solution provided with PM1 and PM2 microplates of Biolog™ Inc. ^82^. The resulting cell suspension was diluted using IF-0 to obtain a 42% T (transmittance) solution by setting it’s OD (600 nm) to 0.376. Further steps were followed as described in the Biolog microplate protocol and PM1 And PM2 were inoculated with the desired *P. agglomerans* R1 genotype. Inoculated plates were then placed in a shaker incubator at 30°C and 200 rpm. Substrate utilization was assessed by measuring the OD of each microplate 24h post inoculation at 590nm. Since certain substrates on the microplate were already coloured, an OD correction was carried out by subtracting the OD value of each well by its respective OD value that was measured before substrate utilization, i.e. immediately after inoculation and before placing the plate in the shaker incubator.

### Determination of maximum respiration rate (μmax)

Maximum growth rate (μmax) was estimated from the first 30 h of Biolog time-course data using a smoothing-spline approach (following ^83^). For each genotype–substrate–replicate combination, a smoothing spline was fitted to OD measurements over time using R’s smooth.spline() function with default parameters. The first derivative of the fitted spline was evaluated across 1,000 evenly spaced time points, and the maximum value was taken as (μmax). For visualization od data, substrates were retained if at least one genotype reached an OD >80 within 30 h, and (μmax) values were calculated independently for each replicate before being averaged across replicates.

### Metabolic pathway predictions

We predicted the metabolic pathways present in *P. agglomerans* R1 (GCF_010667995.1). To this end, we generated four *in silico* knockouts; ΔLPP1, ΔLPP2, ΔLPP1&2 and Δchr, by deleting the LPP1, LPP2 and chromosomal genes from the genome, respectively. These knockouts comprise 3,724, 4,108, 3,578 and 676 genes for ΔLPP1, ΔLPP2, ΔLPP1&2 and Δchr, respectively. Pathway predictions were done for each genotype separately using gapseq version 1.1 (doall command with default parameters) ^84^.

## Acknowledgements

We thank Lisa Hartmann, Hieu Trhong, Mario Santer, Jordi Weijers and Alex Gashanja for critical comments on the manuscript. We thank Rüdiger Stroeh (experimental farm Hohenschulen of Kiel university) for seed samples and Merle Hülter for assistance in the sampling. This research was supported in part through high-performance computing resources available at the Kiel University computing center.

## Author contributions

Conceptualization: N.H. and T.D.; Funding acquisition: T.D.; Plasmid curing: K.R., N.H.; *In vitro* fitness and colonization experiments: K.R., R.S., S.V.; Transcriptomics conceptualization and experiments: K.R., R.S., S.V.; Transcriptomics data analysis: S.V.; Comparative genomics and pangenome analysis: D.R.P.; Omnilog© data retrieval and analysis: S.V., N.H.; Metabolic network reconstruction: D.R.P.; Writing original draft: S.V., K.R., N.H., T.D.; Writing – review & editing, all authors.

## Data availability

SRA of transcriptomics sequencing

SRA of plasmid cured strains sequencing

