## Supplementary information for "Regulatory and metabolic integration accompany plasmid domestication during adaptation to a plant-associated lifestyle"

[1] Institute of General Microbiology, Kiel University, Kiel, Germany
[2] Present address: Institute of Infection, Veterinary and Ecological Sciences, University of Liverpool, Liverpool, UK

[*] These authors contributed equally

Corresponding author: Tal Dagan

### Supplementary material

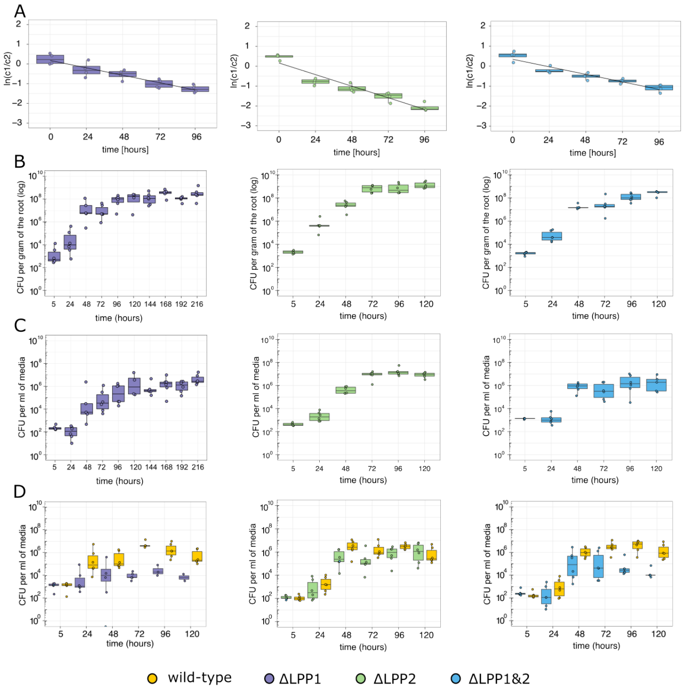

**Figure S1: Fitness assays and wheat root colonization.**

**(A)** *In vitro* competition assays between wild-type *Pantoea* and the indicated plasmid-cured strains in R2A medium at 30 °C. Wild-type cells were labelled with pTW1-mCherry (Wein et al., 2018), and plasmid-cured strains with pRL153-GFP (Tolonen et al., 2007). Plotted are the natural logarithms of the wild-type-to-mutant ratios over time (n=4). **(B, C)** Population dynamics of wild-type and plasmid-cured strains during single-strain wheat colonization experiments, measured as CFU on **(B)** wheat roots and in **(C)** wheat cultivation medium. **(D)** Population dynamics in wheat cultivation medium during competitive root colonization experiments. Wild type, yellow; ΔLPP1, purple; ΔLPP2, green; ΔLPP1&2, blue. Each point in the figure corresponds to an individual plant.

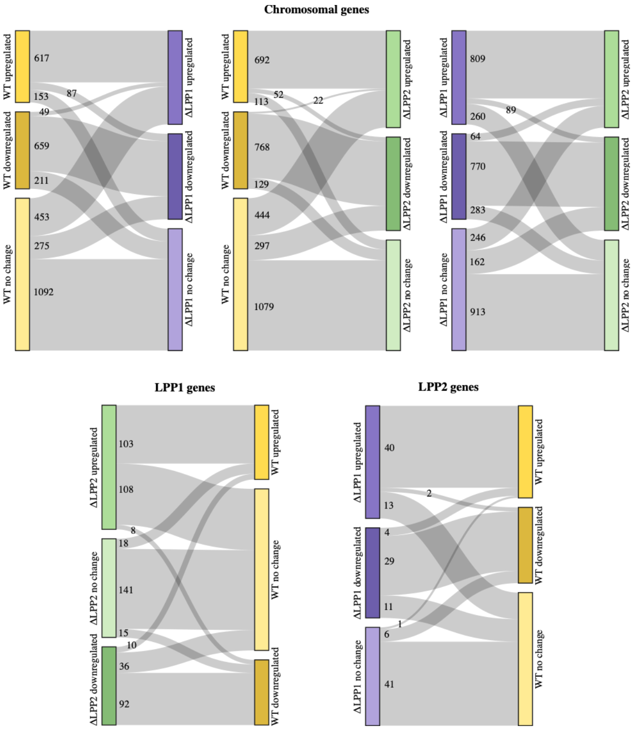

**Figure S2. Comparison of expression patterns between genotypes.**

Sankey diagrams compare genotype-dependent changes in gene expression patterns based on DESeq2 differential-expression analysis. Within each genotype, genes were classified as upregulated, downregulated, or unchanged *in planta* relative to R2A. Each ribbon connects the same set of genes between two genotypes and shows how genes assigned to one expression category in the first genotype were classified in the second genotype. Thus, ribbons remaining within the same category represent genes with conserved expression responses between genotypes, whereas ribbons connecting different categories represent genes whose *in planta* expression response changed depending on genotype. Ribbon width is proportional to the number of genes undergoing each transition, and numbers indicate the corresponding gene counts. Chromosomal genes are compared between wild type (WT) and ΔLPP1, WT and ΔLPP2, and ΔLPP1 and ΔLPP2; LPP1 genes are compared between WT and ΔLPP2, and LPP2 genes between WT and ΔLPP1. WT, yellow; ΔLPP1, purple; ΔLPP2, green.

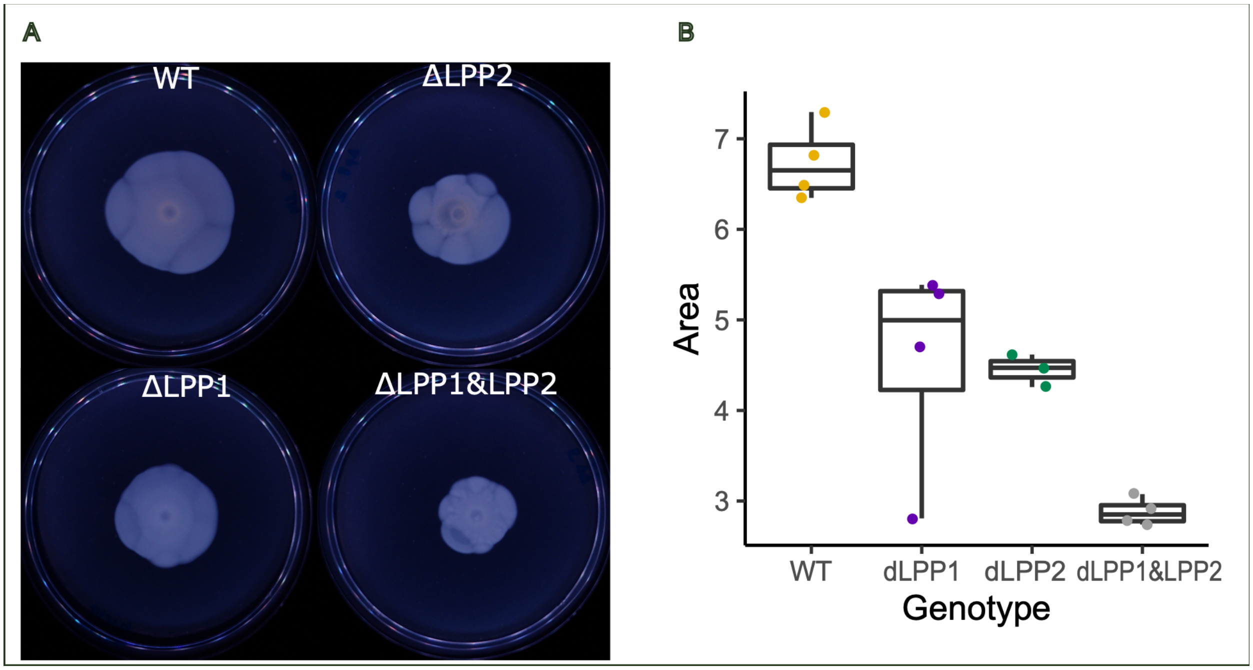

**Figure S3. Comparison of motility among the strains.**

(A) Motility test of wild-type and plasmid-cured strains of *Pantoea* on soft M9+Glucose agar (0.4%) and D) Area covered by colony swarm of each genotype (n = 4)

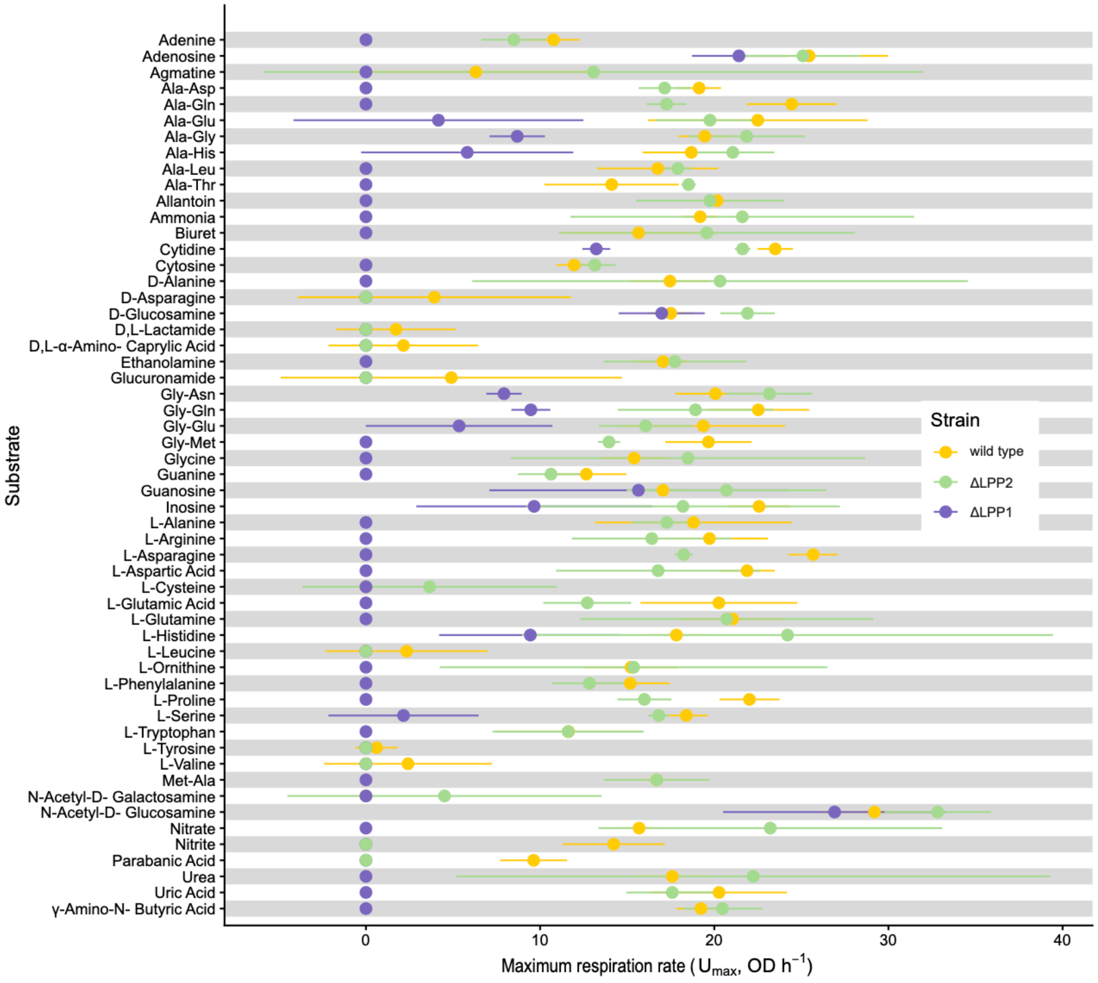

**Figure S4. Utilization of nitrogen sources**

Maximum substrate-dependent respiration rate ($r_{max}$) of wild-type *Pantoea agglomerans* R1 and plasmid-cured derivatives across individual Nitrogen sources, measured using Biolog phenotype microarrays. See Methods for details on the data analysis approach.

**Table S1. Sequence variants detected in wild-type and plasmid-cured strains.**

Sequence variants identified in the wild type (WT), ΔLPP1, ΔLPP2, and ΔLPP1&2 strains are listed together with their frequency and distribution among strains. For each variant, the table provides the genomic location, reference and alternative allele, variant type, affected genomic feature and, where applicable, the predicted coding and amino-acid change. Variant frequencies are reported separately for each genotype, and presence/absence columns indicate the strains in which each variant was detected.

**Table S2. Fitness assays and wheat colonization experiments**

Raw colony-forming unit (CFU) measurements from *in vitro* competition assays, competitive wheat root colonization experiments, and single-strain colonization experiments. The *in vitro* competition sheet contains CFU ml$^{-1}$ measurements for WT and the indicated plasmid-cured strains over time, together with the natural logarithm of the mutant-to-WT CFU ratio. The competitive colonization sheet contains bacterial abundances on wheat roots (CFU g$^{-1}$ root) and in the cultivation medium (CFU ml$^{-1}$), together with the corresponding WT-to-mutant log ratios. The single-colonizer sheet contains CFU measurements for ΔLPP1, ΔLPP2, and ΔLPP1&2 on wheat roots and in the cultivation medium over time. NA indicates measurements for which a ratio or CFU value could not be determined.

**Table S3. Transcriptomics and statistical analysis.**

Transcriptomic analysis of chromosomal, LPP1 and LPP2 genes across genotype and growth-condition comparisons. Genes are listed by locus tag, replicon and protein accession, with the corresponding locus tag in the most recent version of the genome WSSS00000000.2 designated as locus_*tag*_v2. Pairwise differential-expression comparisons as obtained from DEseq2 analysis are reported as log2 fold changes (LFC) and multiple-testing-adjusted P values (padj) for the indicated genotype and condition contrasts. Two-way ANOVA results are additionally provided for comparisons between WT and the indicated mutant genotype across Plant and R2A conditions. For each WT–mutant comparison, false-discovery-rate (FDR)-adjusted P values are reported separately for the main effects of Condition and Genotype, and for the Condition × Genotype interaction. Results are presented on separate worksheets for the chromosome, LPP1 and LPP2 replicons. NA indicates that a value was not available or could not be determined.

**Table S4. Metabolic models.**

Predicted metabolic reactions, associated pathways, and gene assignments for the WT and plasmid-cured strain models. The reactions_genes sheet lists individual metabolic reactions, their associated pathways, the genomic location of the corresponding gene, and the respective locus tag. The cured_models sheet summarizes pathway presence or absence across the different metabolic models, where 1 indicates that the pathway is retained in the model and 0 indicates that it is absent.

**Table S5. Primers used in the study**

**Table S6. Transcript abundance of *Pantoea agglomerans* genes across genotypes and growth conditions.**

Transcript abundance is reported as untransformed transcripts per million (TPM) for individual genes in wild type (WT), ΔLPP1, ΔLPP2, and ΔLPP1 ΔLPP2 strains grown under Plant and R2A conditions. Data are provided on separate worksheets for genes located on the chromosome, LPP1, and LPP2. The chromosome worksheet contains samples from all four genotypes, whereas the LPP1 and LPP2 worksheets contain only genotypes retaining the respective plasmid. For each sample, the genotype, treatment, and replicate are indicated; subsequent columns report TPM values for individual genes.

**Table S7. Pangenome data.**

Each row represents the occurrence of a gene homologous to a gene in *Pantoea agglomerans*. R1 (identified by locus_tag_R1) in the genome indicated by the NCBI Genbank accession column. The corresponding species assignment and replicon location are provided in the inferred species and replicon columns, respectively. The data were obtained from Romero Picazo *et al.* (2025) and were used to estimate gene frequencies across the genus *Pantoea* shown in Figure 3.
